# Leukotriene B_4_ licenses inflammasome activation to enhance skin host defense

**DOI:** 10.1101/2020.02.03.932129

**Authors:** Ana Carolina G Salina, Stephanie Brandt, Nathan Klopfenstein, Amondrea Blackman, Nicole Byers-Glosson, Claudia Brodskyn, Natalia Machado Tavares, Icaro Bonyek Santos Da Silva, Alexandra I de Medeiros, C. Henrique Serezani

**Author notes:** Contributed equally to this work (Ana Carolina G Salina, Stephanie Brandt). Address correspondence: Vanderbilt University Medical Center, Division of Infectious Diseases, Department of Medicine, Nashville, TN, USA. 1161 21^st^ Avenue South, Suite A2310A, MCN, Nashville, TN, 37232.

## Abstract

The initial production of inflammatory mediators dictates host defense as well as tissue injury. Inflammasome activation is a constituent of the inflammatory response by recognizing pathogen and host-derived products and eliciting the production of IL-1β, IL-18 as well as inducing a type of inflammatory cell death termed “pyroptosis”. Leukotriene B_4_ (LTB_4_) is a lipid mediator produced quickly (seconds to minutes) by phagocytes and induces chemotaxis, enhances cytokine/chemokine production, and enhances antimicrobial effector functions. Whether LTB_4_ directly activates the inflammasome is not well understood. Our data show that endogenously produced LTB_4_ is required for the expression of pro-IL-1β *in vivo* and *in vitro* and enhances inflammasome assembly. Furthermore, LTB_4_-mediated Bruton’s tyrosine kinase (BTK) activation is required for inflammasome assembly *in vivo* as well for IL-1β-enhanced skin host defense. Together, these data unveil a new role for LTB_4_ in enhancing the expression and assembly of inflammasome components and suggest that while blocking LTB_4_ actions could be a promising therapeutic strategy to prevent inflammasome-mediated diseases, exogenous LTB_4_ can be used as an adjuvant to boost inflammasome-dependent host defense.

## Introduction

Upon infection, a quick and highly synchronized inflammatory response is mounted to restrict microbial growth and eventually eliminate the pathogen. Therefore, studies focusing on early events could unmask new players involved in the generation and magnitude of the host defense. Inflammasomes are multiprotein, intracellular platforms that detect both pathogen and host-derived products and induce the inflammatory response. The proteins that form the inflammasome consist of upstream sensors that belong to the NOD-like receptor (NLR) family, the adaptor protein, Apoptosis-Associated Speck-Like Protein Containing CARD (ASC), and the downstream effector caspase-1. There are different NLR proteins, including NLRP1, 2, 3, 6, 7, and NLRC4, as well as interferon-inducible protein (AIM2) (1, 2). The different NLRs recognize both distinct and overlapping stressors to elicit the maturation and secretion of the inflammatory cytokines IL-1β and IL-18. Upon cell stimulation with either pathogen - (PAMPs) or damage (DAMPs)-associated molecular patterns, the inflammasome is activated in two sequential steps: 1) transcription of the long forms of IL-1β and IL-18, and 2) the assembly of the inflammasome complex ASC/NLRP and pro-caspase-1and subsequentially autocatalytic cleavage of caspase-1 and processing and secretion of IL-1β and IL-18, as well as production of lipid mediators such as prostaglandins and leukotriene B_4_ (LTB_4_) (3–5). The secretion of these cytokines is followed by a form of programmed cell death called pyroptosis that releases DAMPs and further amplifies the inflammatory response. Increased production of IL-1β and IL-18, along with DAMPs generation, have been heavily implicated in the pathogenesis of a myriad of inflammatory diseases and facilitates antimicrobial activities (1, 6).

*Staphylococcus aureus* skin infection is controlled by the synchronized actions of structural cells (keratinocytes) and skin phagocytes. Inflammasome-dependent IL-1β1β production is required for neutrophil recruitment, abscess formation, and bacterial clearance (7, 8). The mechanisms underlying inflammasome activation and IL-1β1β production during skin infection is not well understood.

*Staphylococcus aureus* skin infection is controlled by the synchronized actions of structural cells (keratinocytes) and skin phagocytes (7, 8). Inflammasome-dependent IL-1β production is required for neutrophil recruitment, abscess formation, and bacterial clearance (9). The mechanisms underlying inflammasome activation and IL-1β production during skin infection is not well understood.

LTB_4_ is a bioactive lipid mediator quickly produced by phagocytes, such as macrophages and neutrophils. LTB_4_ synthesis involves several rate-limiting steps that include activation of phospholipase A_2_ (PLA_2_) and arachidonic acid (AA) release from phospholipids in cellular membranes. Activation of 5-LO and 5-LO activation protein (FLAP) together metabolize AA to LTA_4_, which is converted to LTB_4_ by LTA_4_ hydrolase (LTA_4_H) (10). LTB_4_ binds to two different G protein-coupled receptors (GPCR). LTB_4_ receptor 1 (BLT1) is a high-affinity receptor, and BLT2 has a low affinity to LTB_4_ (11). LTB_4_ via BLT1 enhances inflammatory response by increasing phagocyte chemotaxis, activation of transcription factors required for the production of inflammatory cytokines. LTB_4_ is required for the control of a myriad of pathogens. We and others have shown that LTB_4_NFκB and AP1 and PU.1). LTB_4_ also amplifies pattern recognition response (PRR) by increasing the expression of the TIR adaptor MyD88.NF-κ B and AP-1 and PU.1) (12–15). LTB_4_ also amplifies PRRs response by increasing the expression of the TIR adaptor MyD88 (15). LTB_4_ is required for the control of a myriad of pathogens. We and others have shown that enhances microbial clearance by controlling the generation of reactive oxygen and nitrogen species, as well as antimicrobial peptides *in vivo* (16–19). We also have shown that the treatment of mice with a topical ointment containing LTB_4_ enhances *S. aureus* clearance, production of inflammatory mediators, and abscess formation (20). Whether LTB_4_ enhances skin host defense directly or by indirectly increasing production of proinflammatory mediators remains to be determined. Also, whether influences inflammasome activation is poorly understood. It has been shown that LTB_4_-mediated arthritis severity and increased leishmanicidal activity is impaired in ASC^−/−^ and NLRP3^−/−^ mice (21, 22). However, the mechanisms underlying LTB_4_-mediate inflammasome activation and whether LTB_4_ is a component of the IL-1β-mediated skin host defense remains to be elucidated. Here, using epistatic and gain of function approaches, we are showing that LTB_4_ is required for both first and second signals required for inflammasome activation and identified critical signaling programs required for inflammasome assembly, IL-1β secretion and its consequences in skin host defense.

## Methods

### Animals

Mice were maintained according to National Institutes of Health guidelines for the use of experimental animals with the approval of the Indiana University (protocol #10500) and Vanderbilt University Medical Center (protocol #M1600154) Committees for the Use and Care of Animals. Experiments were performed following the United States Public Health Service Policy on Humane Care and Use of Laboratory Animals and the US Animal Welfare Act. Eighteen-week-old female or male BLT1^−/−^ (B6.129S4-*Ltb4r1^tm1Adl^*/J(23), LysMcre, MMDTR, and strain-matched WT C57BL/6 mice were purchased from Jackson Labs (Bar Harbor, ME USA). EGFP-LysM was donated by Dr. Nadia Carlesso (City of Hope, Duarte, CA, USA), and the pIL1DsRed (donated by Dr. Akiko Takashima, University of Toledo, Toledo, OH, USA (24).

### Primary cell isolation and culture

Murine resident peritoneal macrophages were isolated using ice-cold phosphate-buffered saline (PBS) as previously described (15, 25). To isolate neutrophils, bone marrow from both tibias and femurs were flushed with phosphate-buffered saline (PBS) (20) and neutrophils were negatively isolated using a MACSxpress Neutrophil Isolation Kit, as suggested by the manufacturer (Miltenyi Biotec, Sunnyvale, CA, USA). The media used to culture primary cells was Dulbecco’s Modified Eagle’s Medium (DMEM) supplemented with 5% fetal bovine serum (FBS), 1M HEPES, and 1X antibiotic/antimycotic (GE Healthcare Life Sciences).

### Human phagocyte isolation and culture

Human blood was obtained from healthy donors of Hemoba (Fundação de Hematologia e Homoterapia do Estado da Bahia), Bahia/Brazil. Peripheral blood polymorphonuclear (PMN) cells were isolated by centrifugation using Polymorphprep^TM^ medium according to the manufacturer’s instructions (Axis-Shield, Dundee, UK). Monolayers were harvested and washed with PBS at 4°C, 400g. Cells were cultured in RPMI 1640 medium (Gibco/ThermoFisher Scientific, Waltham, MA, EUA), supplemented with 10% FBS (Gibco), 2 mM L-glutamine, 100 U/ml of penicillin G and 100mg/ml of streptomycin 37°C 5% CO_2_. PMNs were cultured with 100 ng/mL LPS (Sigma-Aldrich) for 2h, washed with PBS and challenged with 10 nM LTB_4_. Culture supernatants were harvested to measure IL-1β release and active caspase-1 by ELISA, as described below.

### Inflammasome activation

Macrophages and neutrophils were challenged with 100ng/mL LPS for 3h, followed by treatment with Nigericin 1μM (1h), Monosodium Urate Crystal (MSU - 10μM), Flagellin 20μg/mL, and Poly(dA:dT)/LyoVec 2μg/mL for 3h. To assess the role of LTB_4_ in amplifying inflammasome activity, cells were treated with 100 nM LTB_4_ or 1 μM BLT1 antagonist U-75302 5 minutes before the addition of the inflammasome activators. The direct effect of LTB_4_ in inflammasome activation was studied in cells treated with LPS as above, followed by LTB_4_ for 3h.

### MRSA skin infection and topical ointment treatment

Mice were infected with MRSA USA300 LAC strain (~3 ×10^6^ colony-forming units [CFU]) *s.c.* in 50μL of PBS as previously shown (26). Lesion and abscess sizes were monitored and determined by affected areas calculated using a standard formula for the area: (A = [π/2] × l × w)(27). The final concentrations of the ointments were as follows: LTB_4_ (33.7 ng– 3.37 × 10^−6^%), U-75302 (0.001%), all in 1g of petroleum jelly (vehicle control). The treatments were applied to the infected area with a clean cotton swab. Mice were treated once a day throughout infection (ranging from 6 hours to 9 days).

### Skin biopsies and bacterial load

Punch biopsies (8 mm) from noninfected (naïve) or infected skin were harvested at different time points and used for determining bacterial counts, cytokine production, RNA extraction, cell isolation, histological analyses, and proximity ligation assay (PLA) staining (28). For bacterial load, skin biopsies were collected on day nine post-infection, processed and homogenized in tryptic soy broth (TSB) medium. Serial dilutions were plated on TSB agar plates, and colonies were incubated at 37ºC, 5% CO_2_ and counted after 18h. Bacterial burdens were normalized to tissue weight and calculated by the following equation: ((CFU/mL plated)*(dilution factor))/(tissue weight in mg). Bacterial burdens in the skin are represented by CFU/mg tissue (26).

### Quantitative real-time PCR (qPCR)

Total RNA was isolated from cultured macrophages or skin biopsies using lysis buffer (RLT; Qiagen, Redwood City, CA, USA). The RT^2^ First Strand Kit reverse transcription system (QIAGEN) was used for cDNA synthesis, and quantitative PCR (qPCR) was performed on a CFX96 Real-Time PCR Detection System (Bio-Rad Laboratories, CA, EUA). Gene-specific primers were purchased from Integrated DNA Technologies (Redwood City, CA, USA). Relative expression was calculated as previously described (29).

### Immunoblotting

Western blots were performed as previously described (13). Cell supernatants and cell lysate were collected, and proteins were precipitated using trichloroacetic acid (TCA). Cell lysate and supernatant were resolved by SDS-PAGE, transferred to a nitrocellulose membrane, and probed with commercially available primary antibodies against caspase-1, IL-1β, ASC (all at 1:1000; Abcam) phosphorylated BTK (Tyr223), total BTK (all at 1:1000; Cell Signaling Technology, Danvers, MA, USA) or β-actin (1:10,000; Sigma-Aldrich, San Luis, Missouri, EUA). Densitometric analysis was performed as described previously (13).

### ELISA assays

Skin biopsy sections were homogenized with a pestle in TNE cell lysis buffer containing 1× protease inhibitor (Sigma-Aldrich, St. Louis, MO, USA) and centrifuged to pellet the cellular debris. The supernatant of cultured macrophages or skin lysate from WT or BLT1^−/−^ mice were also used to detect IL-1β, TNF-α or caspase-1 by ELISA (Biolegend Inc., San Diego, CA, EUA).

### Confocal microscopy and proximity ligation assay (PLA)

PLA was done using Duolink PLA kit (Sigma)in skin sections of naïve and infected WT and BLT1^−/−^ mice or *in vitro* in peritoneal macrophages, and bone marrow neutrophils cultured in 8-well chambered cell culture slides (Corning Inc., Nova York, EUA). *In vitro*, phagocytes were challenged with LPS or GFP-MRSA at MOI 50:1 for 3h and pretreated with 10μM BLT1 antagonist U-75302 or 10nM LTB_4_ for 15 min, followed by inflammasome activation (MSU or nigericin), for 1 hour. For *in vivo* PLA detection, WT mice (treated or not with an ointment containing LTB_4_) or BLT1^−^/^−^ were injected *s.c.* with MRSA for 24h. Skin biopsies were collected for histological analyses and 8μm paraffin-embedded skin sections were used. Cells and skin sections were fixed with 4% formaldehyde for 10 minutes and permeabilized with 0.2% Triton X-100 in tris-buffered saline (TBS) for 10 min. Cells or tissues were then blocked with 1% bovine serum albumin and 10% normal donkey serum in TBS for 1 hour at room temperature and incubated with the indicated primary antibody pairs (ASC versus caspase1 or ASC versus pBTK (Tyr223)) overnight at 4°C. Oligonucleotide-conjugated secondary antibodies (PLA probe MINUS and PLA probe PLUS) against each of the primary antibodies were applied, and ligation and amplification were performed to produce rolling circle products. These products were detected with fluorescently labeled oligonucleotides, and the samples were counterstained with Duolink Mounting Medium with DAPI.

Cells were imaged on a Zeiss LSM 510 confocal microscope with an inverted Axiovert 100 M microscope stand using a C-apochromat 40×/1.2 W corr. Confocal images were taken with identical settings to enable comparison of staining. Z-stacked sections (10 to 22 slices) of the cells were captured in multitrack, and ImageJ software (NIH, Maryland, EUA) was used to reconstruct the images using the Z project plug-in (30, 31). The extent of association between the inflammasome components, as well as their intracellular localization, was quantitated using the JaCoP plug-in for ImageJ (30). The background of the collected images was corrected by the ImageJ rolling ball algorithm plug-in. For each experiment, at least 100 randomly selected cells or 10 field areas were scored.

### Skin biopsy dissociation for flow cytometry

Skin biopsies were digested with collagenase D (Sigma-Aldrich) and processed to obtain a single-cell suspension, as previously described (26). Single cells were stained with the different fluorescent antibodies (as indicated in the figure legends) and DsRed on the BD LSR II flow cytometer (BD Biosciences, San Jose, CA, USA). Analyses were completed using FlowJo analysis software (FlowJo, LLC., Ashland, OR, USA).

### IVIS in vivo imaging

IVIS spectrum/CT (Perkin Elmer, Waltham, MA, EUA) machine was used to image bioluminescence or fluorescence in the mice skin. Mice were anesthetized with isoflurane and imaged while under gas anesthesia. Mice were positioned facing the CCD camera and imaged for the appropriate bioluminescence/fluorescence.

Bioluminescence imaging (BLI) and analysis: The mice were scanned for up to 4 min to allow for bioluminescent signal detection from each mouse. Mice were scanned longitudinally throughout MRSA skin infection. A region was drawn around each infection, and total flux (photons/second) was measured for each infection site. To obtain a background-free total flux signal, the mouse infection region was subtracted from the background region. The background-free total flux was compared to a standard curve to obtain the bacteria amount in each infection. The standard curve was prepared by spotting known bacterial CFU in TSA plates and injected subcutaneously in mice (*in vivo*) and imaged.

DsRed scans and analysis: WT mice that were DsRed served as the autofluorescence background. Mice were scanned, and the Living Image software (Perkin Elmer) was used to unmix the DsRed signal spectrally. A region was drawn around the spectrally unmixed DsRed signal, and total radiant efficiency was measured ([photons/second]/[μW/cm^2^]).

### Statistical analysis

Data analysis were performed in GraphPad Prism software (GraphPad Prism Software Inc., San Diego, CA). The statistical tests used are listed in the figure legends. Briefly, Student’s t-test was used to compare two experimental groups. One-way ANOVA, followed by Tukey multiple comparison correction, was used to compare three or more groups. Two-way ANOVA with repeated measured, followed by Tukey multiple comparison correction, was used to compare infection areas over time between two or more mouse groups. P values < 0.05 were considered significant.

## Results

### LTB_4_/BLT1 axis promotes IL-1β production

LTB_4_ enhances the production of IL-1β during inflammatory conditions. Furthermore, LTB_4_ alone or in combination with cytokines and PAMPs activate different transcription factors that further enhance expression of *Il1b* mRNA expression (32, 33). However, whether LTB_4_ enhances IL-1β abundance by increasing *Il1b* transcripts or IL-1β processing/maturation via effects on inflammasome activation is not known. Here, we measured *Il1b* transcripts and protein by ELISA in skin biopsy homogenates from uninfected and MRSA-infected mice. Skin biopsies from WT mice treated topically with LTB_4_ or the BLT1 antagonist U-75302 (20), as well as Ltb4r1^−/−^ mice collected from uninfected skin and day 1 post-MRSA infection revealed that topical LTB_4_ increased IL-1β and that both BLT1 antagonist treatment and Ltb4r1^−/−^ showed decreased IL-1β protein and *Il1b* mRNA transcripts during infection compared to WT mice **(Figure 1A and B)**. These data suggest that exogenous LTB_4_ (treated topically) enhances inflammasome-dependent IL-1β and endogenously produced LTB_4_ during skin infection is required for optimal IL-1β generation. Next, we aimed to determine whether LTB_4_ could also amplify IL-1β production in human neutrophils. Here, cells were treated with LPS for 3h, followed by LTB_4_ treatment. Our data show that LTB_4_ effects also extends to human neutrophils (**Figure 1F**).

**Figure 1.**
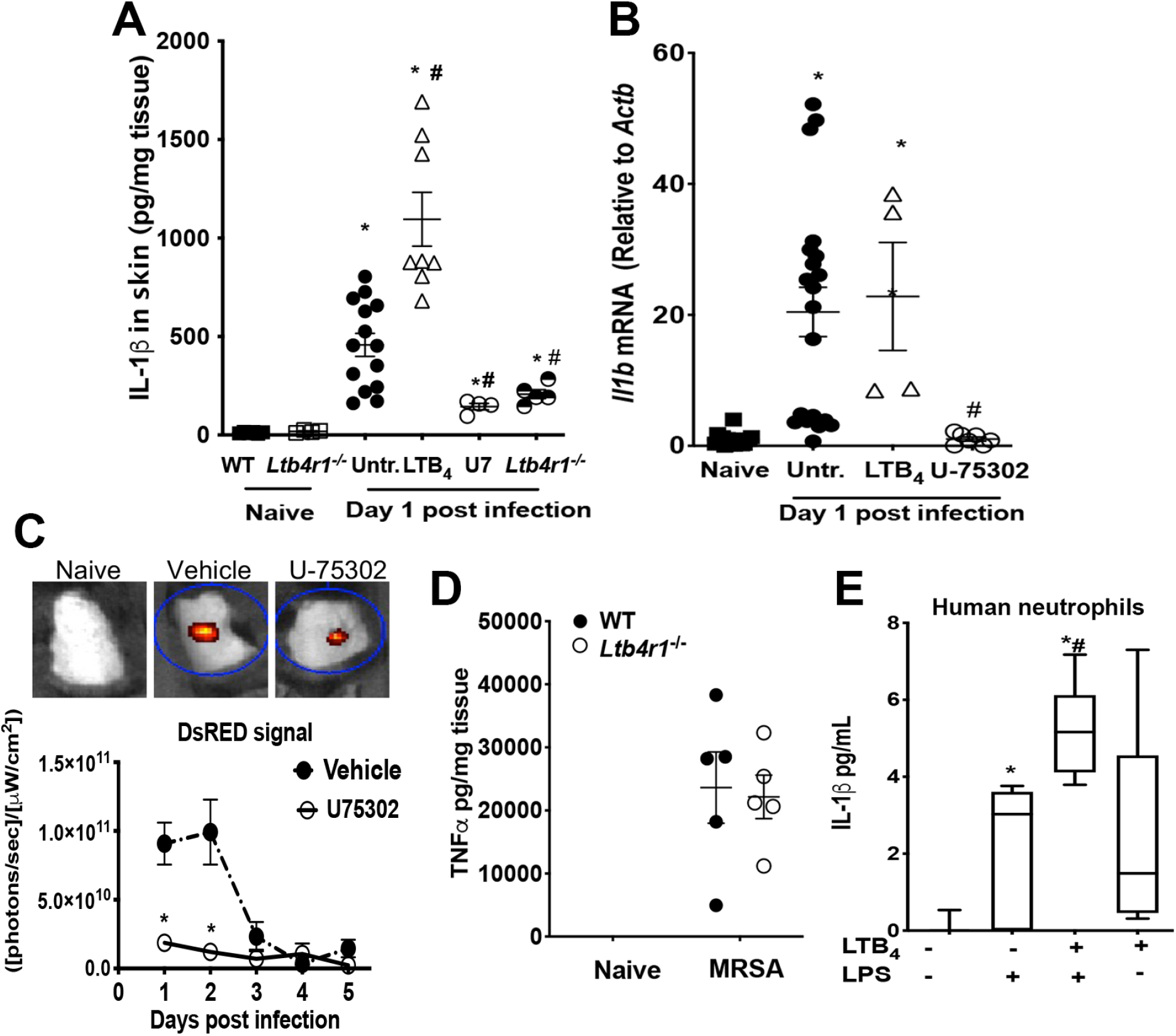
LTB_4_ is required for IL-1β production. **A)** Quantification of IL-1β and **B)** *Il1b* mRNA expression in the skin of C57BL/6, Ltb4r1^−/−^, C57BL/6 treated with a topical ointment containing LTB_4_ or C57BL/6 treated with a topical ointment containing BLT1 antagonist U-75302 24h after MRSA infection. **C)** IVIS scanning of pIL1DsRed mice treated with a topical ointment containing BLT1 antagonist U-75032 or vehicle. TNF-α quantification in **D)** peritoneal macrophages treated with LTB_4_ or BLT1 antagonist before MRSA infection and **E)** skin biopsy from wild-type (WT) or Ltb4r1^−/−^ mice after 24h MRSA infection. **F)** IL-1β quantification from human neutrophils isolated from healthy blood donors treated with LTB_4_ and LPS. Data are the mean ± SEM of 3-10 mice. *p < 0.05 *vs.* naive. #p < 0.05 *vs.* untreated.

IL-1β is produced as a pro-cytokine, which requires gene expression and caspase-1 dependent maturation (34). MRSA skin infection induced expression of *Il1b* **(Figure 1A)**. MRSA-infected mice treated with LTB_4_ ointment did not significantly alter *Il1b* expression; however, BLT1 antagonist U-75302 treatment blunted *Il1b* expression during MRSA skin infection. ELISA data confirmed that IL-1β is produced in the skin of mice during MRSA skin infection and that LTB_4_ treatment increased levels of IL-1β **(Figure 1B)**. Increased levels of IL-1β despite no differences in gene expression suggest that LTB_4_ may be involved in promoting IL-1β processing.

To determine whether LTB_4_ influences the kinetics of IL-1β during MRSA skin infection, we employed the IL-1β reporter mouse pIL1DsRED using IVIS *in vivo* imaging technology. The pIL1DsRED mice were infected with MRSA and treated with a topical ointment containing BLT1 antagonist U-75032 or vehicle control. Mice were imaged by IVIS for DsRed signal everyday post-infection for 5 days **(Figure 1C)**. DsRed signal was highest at days 1 and 2 post-infection and decreased by day 3 of infection **(Figure 1C)**. This suggests that IL-1β is expressed early during MRSA skin infection. BLT1 antagonist treatment showed minimal expression of DsRed at all points measured during infection, indicating that LTB_4_ was necessary for IL-1β expression during MRSA skin infection.

Furthermore, the specific effect of LTB_4_/BLT1 in *Il1b* mRNA expression was evidenced by the fact that neither exogenous LTB_4_ or BLT1 antagonist influences TNF-α expression and secretion in macrophages (**Supplementary Fig. 1 A and B**). Moreover, we did not detect differences in TNF-α production after infection in the skin biopsy from Ltb4r1^−/−^ mice (**Figure 1E**). Pharmacological and genetic BLT1 inhibition showed lower levels of IL-1β at day 1 post-MRSA skin infection than WT mice.

Next, we aimed to identify in which cells BLT1 is necessary for IL-1β production during MRSA skin infection. The pIL1DsRED mice were infected with MRSA and treated with or without BLT1 antagonist U-75302. BLT1 antagonist treatment did not alter the overall percent of macrophages (F4/80+ cells) in the skin during MRSA skin infection **(Supplementary Figure 2A)**; however, the percent of macrophages expressing *Il1b* (DsRed^+^) was significantly lower in the skin of BLT1 antagonist-treated mice than macrophages in the skin of untreated mice **(Supplementary Figure 2B)**. Not only the percent of total F4/80^+^ macrophages expressing DsRed was reduced with BLT1 antagonist treatment, but also the mean fluorescence intensity (MFI) of DsRed was reduced in cells expressing DsRed **(Supplementary Figure 2C)**. BLT1 antagonism decreased the number of global Ly6G^+^ neutrophils but did not reduce the numbers of DsRed expressing PMNs **(Supplementary Figure 2D)**. Importantly, the MFI of DsRed was lower in mice treated with BLT1 antagonist **(Supplementary Figure 2E and F),** indicating that BLT1 activation was not required for PMN migration to the infected skin, but rather, necessary for PMN IL-1β1β expression.

### LTB_4_ enhances NLRP3 inflammasome activity

We then tested whether LTB_4_ controlled IL-1β processing through inflammasome assembly and caspase-1 activation. To determine inflammasome assembly, bone marrow-derived neutrophils were imaged by fluorescence microscopy to determine associations of NLRP3 and ASC using PLA, and the close proximity of these proteins resulted in red fluorescence. Neutrophils with no treatment did not show interactions between NLRP3 and ASC **(Figure 2A)**. Neutrophils co-cultured with MRSA induced minimal, but significant, inflammasome assembly. Nigericin-induced inflammasome activation resulted in high NLRP3/ASC association compared to vehicle only. While LTB_4_ treatment further promoted, the BLT1 antagonist U-75302 treatment inhibited inflammasome assembly **(Figure 2A)**. To confirm these data, we used another inflammasome activator, MSU in peritoneal macrophages from WT or BLT1^−/−^ mice; BLT1^−/−^ macrophages treated with LPS plus MSU induced less inflammasome assembly than WT macrophages (**Figure 2B**).

**Figure 2.**
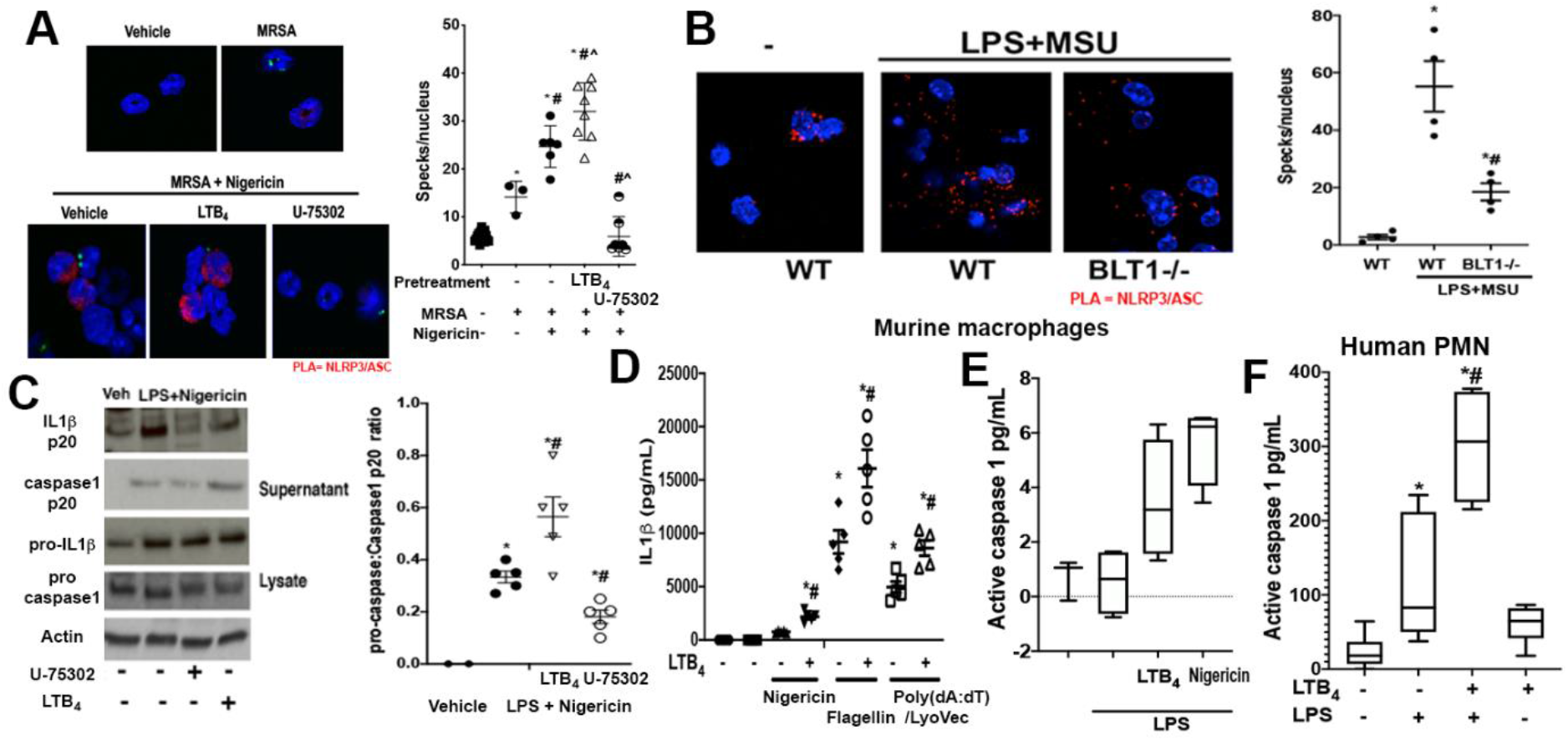
LTB_4_ enhanced inflammasome assembly. Proximity-ligation assay (PLA) was used to detect associations between NLRP3 and ASC represented by red signal of **A)** Bone marrow neutrophils pretreated with 10nM LTB_4_ for 5 minutes, 10μM U-75302 for 30 minutes and cultured with or without GFP-MRSA for 3h followed by 1μM nigericin for 1h and **B)** Peritoneal macrophages from WT or BLT1^−/−^ mice treated with 100ng/mL LPS for 3h followed by 10 μM MSU for 1h. **C)** Cleavage of IL-1β and caspase 1 evaluation by immunoblotting of macrophages treated with 100ng/mL LPS and 1μM nigericin in the presence of LTB_4_ and BLT1 antagonist. **D)** Macrophages were stimulated with LTB_4_ plus different inflammasome activators (Nigericin 1μM, flagellin 20μg/mL and Poly(dA:dT)/LyoVec 2μg/mL) and IL-1β was determined. Murine macrophages **(E)** and Human PMN **(F)** were treated with 100ng/mL LPS for 3h followed by 10nM LTB_4_ and 1μM nigericin and active caspase-1 was measured on the supernatant. Quantification of the intensity of the red signal and western blotting bands was measured by ImageJ analysis. Data are mean ± SEM at least 10 cells from 3-6 mice *p < 0.05 *vs.* untreated cells or LPS + MSU. #p < 0.05 *vs.* MRSA only or treatment with LTB_4_. ^p < 0.05 *vs.* MRSA + nigericin.

These findings were confirmed as we observed decreased active caspase-1 and mature IL-1β in the supernatant of macrophages treated with BLT1 antagonist and increased IL-1β in LTB_4_ treated cells **(Figure 2C)**. Next, we aimed to investigate whether LTB _4_LTB_4_ effects are restricted to NLRP3 or also enhance the activation of the AIM2 or NLRC4 inflammasome. Macrophages were challenged with LPS, followed by LTB_4_ and the AIM2 activator (Poly(dA:dT)/LyoVec) or the NLRC4 agonist (flagellin). In all circumstances, LTB_4_ further increased IL-1β abundance when macrophages were challenged with these other inflammasome activators (**Figure 2D**). Also, murine macrophages and human neutrophils treated with LTB_4_ alone showed increased active caspase-1 when compared to cells treated with inflammasome activators. Then, we studied whether LTB_4_ itself drives the second signal to activate inflammasome and induce IL-1β secretion. Here, macrophages were treated with LPS as above, followed by LTB_4_ for 3h. We detected IL-1β increased in the supernatant of macrophages treated with LPS plus LTB_4_ when compared to LPS or LTB_4_ alone. Furthermore, increased IL-1β in the supernatant correlated with enhanced levels of active caspase-1 (**Figure 2E**). That LTB_4_ itself enhances inflammasome activation was also evidenced in human neutrophils **(Figure 2F)**. These results suggest that LTB_4_/BLT1 is necessary for *Il1b* expression and that LTB_4_ may be an essential mediator in regulating inflammasome assembly and activation, allowing for mature IL-1β production.

Next, we were poised to test the role of LTB_4_ in inflammasome activation *in vivo*. Initially, we challenged WT and BLT1^−/−^ mice with MSU, *i.p*, and 24h later we detected lower IL-1β in the peritoneal lavage of BLT1^−/−^ mice than WT mice challenged with MSU (**Figure 3A)**. We then determined whether LTB_4_ production is required for inflammasome activation during MRSA skin infection. WT mice were challenged with MRSA *s.c.*, followed by topical treatment with LTB_4_, as we have previously shown (20). Our data show that while MRSA skin infection increases the abundance of both IL-1β and caspase-1, topical LTB_4_ further enhances IL-1 β secretion and caspase-1 activation in skin lysates **(Figure 3B)**. Importantly, we confirmed the role of BLT1 in inflammasome activation *in vivo* in skin sections stained for the association between ASC/caspase-1 using PLA. Our data showed decreased ASC/caspase-1 interaction in areas near the abscess in MRSA-infected BLT1^−/−^ mice when compared to infected WT mice **(Figure 3C)**.

**Figure 3.**
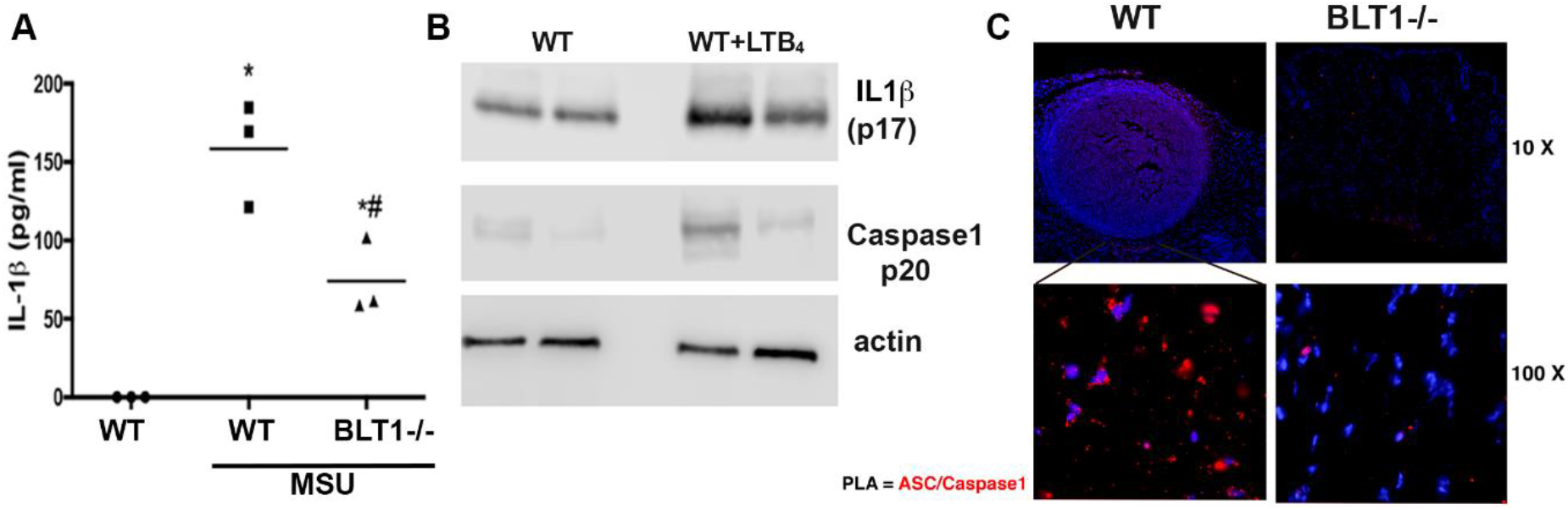
LTB_4_/BLT1 axis is important to inflammasome assembly and IL-1β production. **A)** Quantification of IL-1β production of WT and BLT1^−/−^ mice inoculated with 1 ug/mL MSU or PBS. **B)** IL-1β and caspase-1 activation measured by western blotting of from MRSA-infected WT mice treated with ointment containing vehicle control or LTB_4_ and. **C)** PLA was used to detect associations between caspase-1 and ASC represented by the red signal in the MRSA infected skin of WT and BLT1^−/−^ mice. Data are the mean ± SEM of 3-6 mice. * p < 0.05 *vs.* WT+PBS. #p < 0.05 *vs.* WT+MSU.

Together, these data show that endogenously produced LTB_4_ during inflammatory response or skin infection is required for both first and second signals for inflammasome activation.

### LTB_4_/BLT1 axis enhances BTK-mediated inflammasome activation

Inflammasomes are controlled by a variety of post-translational modifications, including phosphorylation (35). Here, we tested the role of different kinases known to be both activated by LTB_4_ and, involved in inflammasome activation (35). Macrophages were challenged with LPS, followed by the indicated inhibitors (see legend), LTB_4_, and then nigericin. Our data show that only the BTK inhibitor (ibrutinib), but not PI3K, PKCδ, and Syk inhibitors prevented LTB_4_-mediated enhanced IL-1β production, **(Figure 4A)**. The role of BTK, but not other kinases in LTB_4_-mediated inflammasome activation was further confirmed by detecting active caspase-1 in the cell supernatant by immunoblotting **(Figure 4B)**.

**Figure 4.**
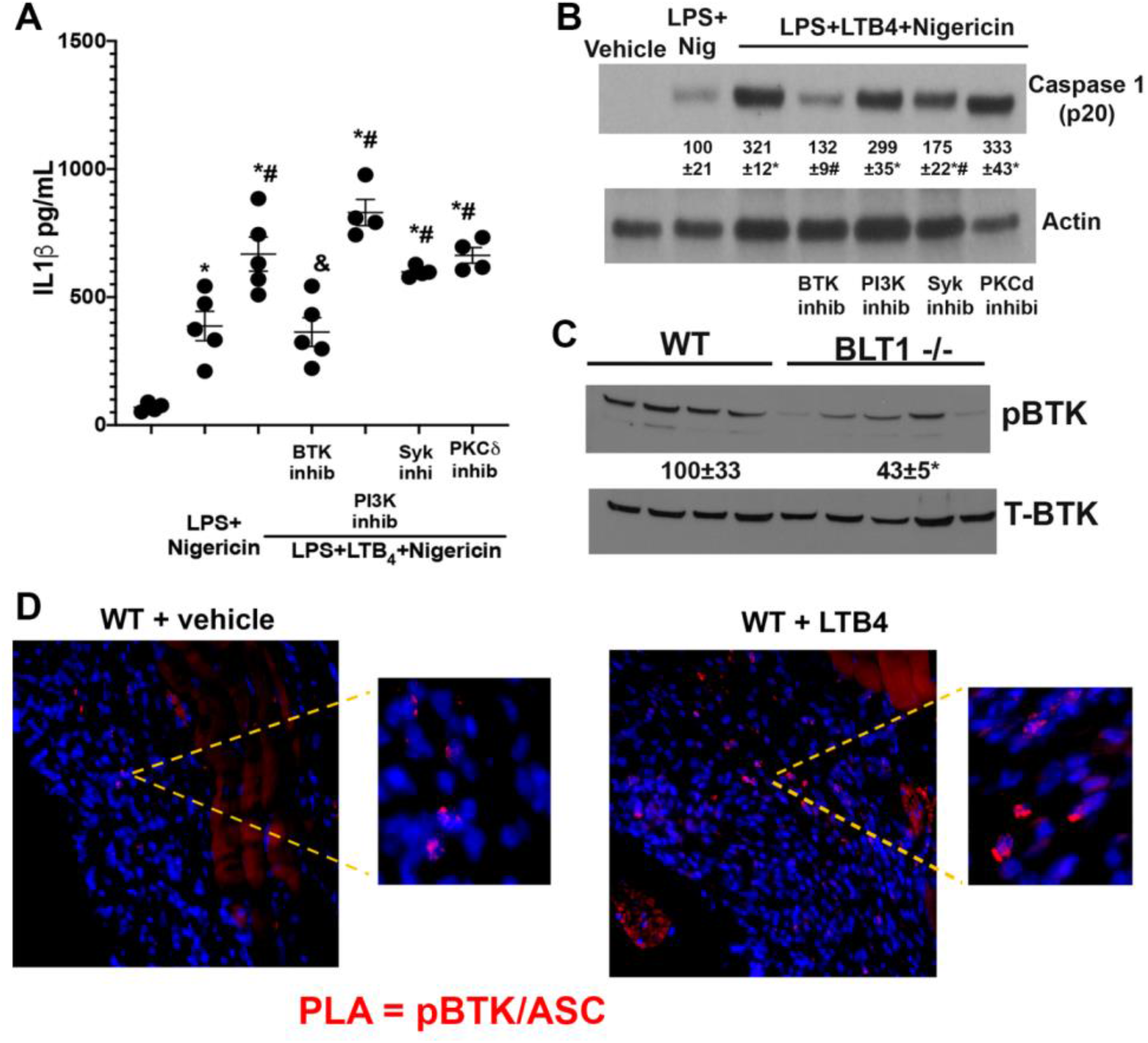
LTB_4_-induced BTK phosphorylation is required for inflammasome assembly. Macrophages were stimulated with 100 ng/mL LPS for 3h, followed by treatment with BTK inhibitor (Ibrutinib), PI3K inhibitor (wortmanin), Syk inhibitor (R406) and PKCδ (Rottlerin) for 20 min and 10nM LTB_4_ for 10 min and 1uM nigericin treatment for 1h. **A)** IL-1β quantification by ELISA. **B)** Western blotting analysis of caspase-1 activation. The numbers represent mean densitometric analysis of the bands shown from 3 independent experiments. **C)** WT and BLT1^−/−^ mice were infected with MRSA for 24 h and skin lysates from WT or BLT1^−/−^ mice were subjected to immunoblotting, followed by detection of total and phosphorylated BTK (Tyr223). **D)** PLA of pBTK (Tyr223) and ASC in the infected skin of WT topically with vehicle control or LTB_4_. Data are the mean ± SEM of 5 mice from at least 2 individual experiments. *p < 0.05 *vs.* untreated cells. #p < 0.05 *vs.* LPS+Nigericin. & p < 0.05 *vs.* LPS+LTB_4_+Nigericin.

Whether LTB_4_ activates BTK remains to be determined. Here, we assessed the role of BLT1 in BTK phosphorylation and expression during MRSA skin infection. Our data clearly show that BTK is activated during MRSA infection and that LTB_4_/BLT1 signaling is required for optimal BTK phosphorylation in the skin (**Figure 4C**). Given the role of LTB_4_ in BTK actions, we aimed to identify whether LTB_4_ would enhance BTK and ASC association *in vivo*. Using PLA to show protein interaction, our data unveiled that MRSA skin infection leads to enhanced phosphorylated BTK and ASC interaction, and importantly, topical LTB_4_ treatment further increased ASC/BTK association (**Figure 4D**). In summary, we are showing that LTB_4_ is required for different steps of inflammasome activation by increasing transcriptional programs, as well as increasing phosphorylation of ASC and increasing speck formation.

### Crosstalk between LTB_4_ and IL-1β to enhance skin host defense

To demonstrate whether LTB_4_ is also involved in IL-1β-mediated skin host defense, we performed “add-back” experiments, by injecting WT and BLT1^−/−^ mice with recombinant IL-1β *s.c* at the moment of infection. Our data show that IL-1β increases bacterial load and decreases lesion size in BLT1^−/−^ mice (**Figure 5 A**). Interestingly, IL-1β decreases bacterial load in WT mice but increases lesion size (as evidenced by elevated inflammatory response) (**Figure 5 B**). Together, these data show that LTB_4_ production is a crucial component of IL-1β effects during skin bacterial clearance.

**Figure 5.**
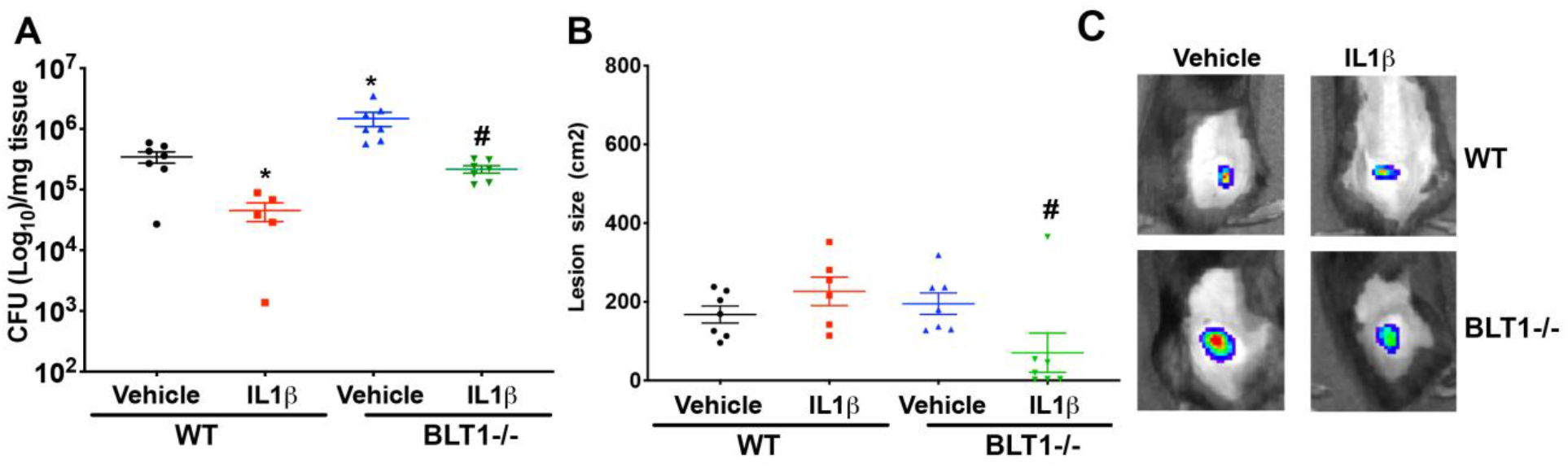
The addition of IL-1β restores deficient skin host defenses in BLT1^−/−^ mice. WT or BLT1^−/−^ mice were injected *s.c.* with MRSA with or without 50ng/mL IL-1β for 24h. **A)** Colony-forming units (CFU) were measured from the skin biopsies of animals on day nine. **B)** The skin size lesion determined on day 9. **C)** Representative bioluminescence MRSA quantification in mice inoculated in the presence or absence of IL-1β using IVIS Spectrum. Data are the mean ±SEM of 5-7 from 2 individual experiments. * p<0,05 *vs.* WT. #p < 0.05 *vs.* BLT1^−/−^.

## Discussion

Inflammasome activation is a critical component of both host defense and inflammatory responses. Multiple regulatory programs enhance inflammasome activation, including enhancing transcriptional programs, ROS production, acetylation, ubiquitination and phosphorylation (35). However, the signals that trigger upstream kinase activation and amplifies inflammasome assembly remains to be fully determined. Here, we are moving the field forward by identifying a regulatory program that amplifies both first and second signal required for optimal inflammasome activation and IL-1β production. Our data show that 1) genetic and pharmacologic LTB_4_/BLT1 actions promotes macrophage pro-*Il1b* mRNA expression *in vivo* and *in vitro*; 2) exogenous LTB_4_ enhances inflammasome-dependent IL-1β secretion; 3) Both exogenous and endogenous LTB_4_ enhances active caspase-1 and mature IL-1β in the supernatant of macrophages and neutrophils from both mice and humans; 4) LTB_4_ increases NLRP3 inflammasome assembly in a manner dependent on BTK phosphorylation; and 5) LTB_4_/BLT1 signaling is required for IL-1β-dependent MRSA skin host defense.

LTB_4_ participates in the initiation, potency, maintenance, and endurance of the inflammatory response by regulating the actions of different immune cells (36). LTB_4_ is a potent inflammatory mediator that has long known to be a robust neutrophil chemoattractant(37, 38). More recently, new roles for LTB_4_ in phagocyte biology has emerged. We and others have shown that LTB_4_ is required for phagocytosis and antimicrobial effector functions by increasing reactive oxygen and nitrogen species secretion and defensin production during infection by gram-positive and negative bacteria, protozoan parasites, and viruses (16–19, 39). Although LTB_4_ is an essential homeostatic mediator of the inflammatory response, high and sustained LTB_4_ production has been associated with inflammatory diseases and metabolic dysfunctions (36, 40), supporting the idea that abundant and chronic circulating LTB_4_ levels are harmful to the host. Moreover, exaggerated LTB_4_ is responsible for an aberrant TNF-α production that drives susceptibility to mycobacterial infection in a zebrafish model. Also, a gain-of-function single-nucleotide polymorphism in the LTA_4_H gene is associated with susceptibility to tuberculosis in humans (41, 42). Furthermore, we also have shown that during reduced skin infection in diabetic mice is characterized by an overwhelming inflammatory response, characterized by abundant LTB_4_ and IL-1β production and that blocking BLT1, but not BLT2 restores IL-1β to basal levels in the infected skin (43). We also have shown that blocking LTB_4_ synthesis shows beneficial effects in a model of CLP-induced sepsis in diabetic mice by decreasing IL-1β IL-1β production and MyD88 expression (29). Here, we are further showing that blocking BLT1 in NLRP3 inflammasome-activated macrophage from diabetic mice inhibited caspase-1 activation (**Supplementary Figure 3**).

Although we and others have shown that IL-1β participates in detrimental LTB_4_ responses, whether LTB_4_ differently regulates the first and/or the second signal required for inflammasome-dependent IL-1β maturation is unknown. LTB_4_-induced IL-1β production and NLRP3 inflammasome activation has been shown during L*eishmania amazonensis* infection. (44) and in a model of arthritis (21, 22). However, to date, no mechanistic studies have been done to address this critical question.

We and others have shown that LTB_4_ controls the expression and actions of different PRRs, oxidant generation, protease release, and activation of inflammatory transcription factors (13, 14, 31–33, 45). LTB_4_ increases NF-κB activation that could, in turn, enhance IL-1β, NLRP3, and caspase-1 expression. Here, using a reporter mouse that detects IL-1β (3, 13). Here, using a reporter mouse that detects expression (IL1bDsRed), we are showing that BLT1 antagonism inhibits *Il1b* transcription in both macrophages and neutrophils at the peak of expression post-infection. This effect was restricted to IL-1β since we did not witness an effect of topical LTB_4_ or BLT1 antagonist in TNF-α production in macrophages during MRSA skin infection. Since NF-κB controls the expression of both IL-1β and TNF-α, the specific mechanisms underlying the effects of LTB_4_ in IL-1β actions remain to be determined.

We and others have shown that the LTB_4_/BLT1 axis either directly induces or further amplifies the activation of different kinases, shaping cellular responses in different environments (45–47). The inflammasome is activated by signals derived from pathogen products and host-derived molecules (DAMPs). Here, we are expanding the list of endogenous molecules that activate inflammasome, e.g., LTB_4_. However, the novelty of our work stems from the fact that LTB_4_ is not secreted by dying cells. LTB_4_ is produced quickly (seconds to minutes) after cell activation and therefore, could further amplify the signals required for inflammasome assembly. Here, we showed that LTB_4_ amplifies the effects of different NLRP3 agonists as well as induce inflammasome assembly directly. We then determined the events by which LTB_4_ enhances NLRP3 inflammasome assembly.

Given the role of BLT1 in activating a myriad of kinases, including ones known to enhance inflammasome activation (48–50), we tested the role of BLT1 in the NLRP3 inflammasome assembly. Our data show that BLT1 activation is required for both *in vivo* and *in vitro* inflammasome activation in a variety of systems and activators, LTB_4_/BLT1 is required for MSU, nigericin and MRSA-induced inflammasome activation, which shows the broad effects of LTB_4_ in inflammasome activation. The role of eicosanoids in inflammasome activation remains inconclusive. While we and others have shown that prostaglandin E_2_ (PGE_2_) enhances inflammasome activation in a model of scorpion venom, Zoccal *et al*, have shown that PGE_2_/Eprostanoid receptor (EP) enhances PKA-dependent NLRP3 phosphorylation in macrophages. PGE_2_/PKA signaling is dictated by Gas-dependent cAMP production (3, 51).

We and others have shown that BLT1 is coupled to both G_αi_ and G_αq_ that inhibits cAMP and increases Ca^2+^ mobilization, respectively(15, 52, 53). Importantly, GPCRs that are coupled to either G_αi_ or G_αq_ are also known to activate the inflammasome, by directly modulating the abundance of second messengers, or by influencing kinase-mediated inflammasome association. Although we did not investigate the role of different G proteins in BLT1-mediated inflammasome activation and the role of downstream effectors, the fact that neither PI3K or PKC-δ inhibition prevented LTB_4_-mediated inflammasome activation, suggests that G_αi_-dependent decreased cAMP might be allowing inflammasome assembly. Whether cAMP influences BTK activation remains to be determined.

Here, we provided evidence that LTB_4_/BLT1/BTK phosphorylation is required for NLRP3/ASC assembly in phagocytes. Our data show that LTB_4_-mediated BTK1 phosphorylation is needed for the NLRP3/ASC association during inflammasome activation as well as during MRSA skin infection. BTK is a tyrosine kinase known to be necessary for B cell receptor and FcR signaling(54). More recently, the role of BTK in phagocytes has emerged. BTK activation is required for TLR signaling, phagocyte recruitment to the inflammatory site, and macrophage maturation (55, 56). BTK is phosphorylated in two tyrosine sites (Y223 and Y551), and Syk and Lyn are known to be upstream to BTK in B cells (57). Whether these kinases are involved in BTK actions in macrophages is unknown. Here, we showed that Syk inhibitor R406 did not prevent LTB_4_-mediated inflammasome, which suggests that Syk does not influence BTK actions in inflammasome actions. BTK classically activates PLC, PI3K/Akt, and NF-κB, which could, in turn, regulate the first signal (NF-κB-dependent expression of inflammasome components) as well as the second signal (PI3K) and therefore, increasing NLRP3 phosphorylation and favoring inflammasome assembly. However, PI3K inhibition does not prevent LTB_4_ effects on inflammasome assembly. These data suggest that LTB_4_ enhances inflammasome assembly by increasing the association between phosphorylated BTK with ASC. Future studies will be needed to determine the events upstream to BTK activation, as well as the ASC phosphorylation sites activated by BTK.

IL-1β is crucial for the control of *S. aureus* skin infection by controlling neutrophil migration to the site of infection and abscess formation(58). We have demonstrated that LTB_4_ also enhances *S. aureus* clearance in the skin by increasing neutrophil migration and abscess formation (20). We and others have shown that LTB_4_ enhances IL-1β, but here, we are providing further evidence that both IL-1β and LTB_4_ are part of a positive feedback loop required for optimal host defense. Importantly, it has been shown that LTB_4_ is produced as part of the inflammasome activation, an event called eicosanoid storm(59). In preliminary experiments, when we infected the skin of caspase-1 and NLRP3 deficient mice, we did not observe a reduction in LTB_4_ levels (not shown), suggesting that LTB_4_ is both an initial key component of the IL-1β-dependent skin bacterial clearance and that LTB_4_ is also part of a continuous positive feedback loop to protect the skin from infection.

In summary, our data show that the prevention of exaggerated LTB_4_/BLT1 actions is a promising and potent therapeutic strategy to prevent overwhelming inflammatory response by inhibiting both transcriptional programs involved in the expression of inflammasome components as well as preventing the intracellular signals required for inflammasome assembly. On the other hand, the treatment with exogenous LTB_4_ is a promising candidate to increase both antimicrobial effector functions and inflammatory response in hard to treat infections.

## Acknowledgments

We thank Bethany Moore (University of Michigan, Ann Arbor, Michigan, USA) for providing the MRSA USA300 LAC strain and Roger Plaut (Food and Drug Administration, Silver Spring, Maryland, USA) for providing the bioluminescent USA300 (NRS384 lux) MRSA strain. This work was supported by NIH grants R01HL124159-01 and RAI149207A (to CHS) and T32AI060519 (to SLB).

## References

1. Broz P. Recognition of intracellular bacteria by inflammasomes. Bacteria and Intracellularity. 2019:287.

2. Sharma D, and Kanneganti T-D. The cell biology of inflammasomes: Mechanisms of inflammasome activation and regulation. Journal of Cell Biology. 2016;213(6):617–29.

3. Zoccal KF, Sorgi CA, Hori JI, Paula-Silva FW, Arantes EC, Serezani CH, et al. Opposing roles of LTB 4 and PGE 2 in regulating the inflammasome-dependent scorpion venom-induced mortality. Nature communications. 2016;7(1):1–13.

4. von Moltke J, Trinidad NJ, Moayeri M, Kintzer AF, Wang SB, van Rooijen N, et al. Rapid induction of inflammatory lipid mediators by the inflammasome in vivo. Nature. 2012;490(7418):107–11.

5. Rathinam VA, and Fitzgerald KA. Inflammasome complexes: emerging mechanisms and effector functions. Cell. 2016;165(4):792–800.

6. Kesavardhana S, and Kanneganti T-D. Mechanisms governing inflammasome activation, assembly and pyroptosis induction. International Immunology. 2017;29(5):201–10.

7. Abtin A, Jain R, Mitchell AJ, Roediger B, Brzoska AJ, Tikoo S, et al. Perivascular macrophages mediate neutrophil recruitment during bacterial skin infection. Nature immunology. 2014;15(1):45.

8. Brandt SL, Putnam NE, Cassat JE, and Serezani CH. Innate Immunity to Staphylococcus aureus: Evolving Paradigms in Soft Tissue and Invasive Infections. The Journal of Immunology. 2018;200(12):3871–80.

9. Miller LS, Pietras EM, Uricchio LH, Hirano K, Rao S, Lin H, et al. Inflammasome-mediated production of IL-1β is required for neutrophil recruitment against Staphylococcus aureus in vivo. The Journal of Immunology. 2007;179(10):6933–42.

10. Werz O. 5-lipoxygenase: cellular biology and molecular pharmacology. Current Drug Targets-Inflammation & Allergy. 2002;1(1):23–44.

11. Yokomizo T. Two distinct leukotriene B4 receptors, BLT1 and BLT2. The Journal of Biochemistry. 2015;157(2):65–71.

12. Pasparakis M, Haase I, and Nestle FO. Mechanisms regulating skin immunity and inflammation. Nature reviews immunology. 2014;14(5):289–301.

13. Serezani CH, Lewis C, Jancar S, and Peters-Golden M. Leukotriene B 4 amplifies NF-κB activation in mouse macrophages by reducing SOCS1 inhibition of MyD88 expression. The Journal of clinical investigation. 2011;121(2):671–82.

14. Serezani CH, Kane S, Collins L, Morato-Marques M, Osterholzer JJ, and Peters-Golden M. Macrophage dectin-1 expression is controlled by leukotriene B4 via a GM-CSF/PU. 1 axis. The Journal of Immunology. 2012;189(2):906–15.

15. Wang Z, Filgueiras LR, Wang S, Serezani APM, Peters-Golden M, Jancar S, et al. Leukotriene B4 enhances the generation of proinflammatory microRNAs to promote MyD88-dependent macrophage activation. The Journal of Immunology. 2014;192(5):2349–56.

16. Gaudreault É, and Gosselin J. Leukotriene B4 induces release of antimicrobial peptides in lungs of virally infected mice. The Journal of Immunology. 2008;180(9):6211–21.

17. Wan M, Sabirsh A, Wetterholm A, Agerberth B, and Haeggstrom JZ. Leukotriene B4 triggers release of the cathelicidin LL-37 from human neutrophils: novel lipid-peptide interactions in innate immune responses. The FASEB Journal. 2007;21(11):2897–905.

18. Serezani CH, Aronoff DM, Jancar S, and Peters‐Golden M. Leukotriene B4 mediates p47phox phosphorylation and membrane translocation in polyunsaturated fatty acid‐stimulated neutrophils. Journal of leukocyte biology. 2005;78(4):976–84.

19. Wymann MP, von Tscharner V, Deranleau DA, and Baggiolini M. The onset of the respiratory burst in human neutrophils. Real-time studies of H2O2 formation reveal a rapid agonist-induced transduction process. Journal of Biological Chemistry. 1987;262(25):12048–53.

20. Brandt SL, Klopfenstein N, Wang S, Winfree S, McCarthy BP, Territo PR, et al. Macrophage-derived LTB4 promotes abscess formation and clearance of Staphylococcus aureus skin infection in mice. PLoS pathogens. 2018;14(8):e1007244.

21. Chen M, Lam BK, Kanaoka Y, Nigrovic PA, Audoly LP, Austen KF, et al. Neutrophil-derived leukotriene B4 is required for inflammatory arthritis. The Journal of experimental medicine. 2006;203(4):837–42.

22. Tavares NM, Araújo-Santos T, Afonso L, Nogueira PM, Lopes UG, Soares RP, et al. Understanding the mechanisms controlling Leishmania amazonensis infection in vitro: the role of LTB4 derived from human neutrophils. The Journal of infectious diseases. 2014;210(4):656–66.

23. Tager AM, Bromley SK, Medoff BD, Islam SA, Bercury SD, Friedrich EB, et al. Leukotriene B 4 receptor BLT1 mediates early effector T cell recruitment. Nature immunology. 2003;4(10):982–90.

24. Matsushima H, Ogawa Y, Miyazaki T, Tanaka H, Nishibu A, and Takashima A. Intravital imaging of IL-1beta production in skin. J Invest Dermatol. 2010;130(6):1571–80.

25. Alvarez ARP, Glosson-Byers N, Brandt S, Wang S, Wong H, Sturgeon S, et al. SOCS1 is a negative regulator of metabolic reprogramming during sepsis. JCI insight. 2017;2(13).

26. Dejani NN, Brandt SL, Piñeros A, Glosson-Byers NL, Wang S, Son YM, et al. Topical Prostaglandin E Analog Restores Defective Dendritic Cell–Mediated Th17 Host Defense Against Methicillin-Resistant Staphylococcus Aureus in the Skin of Diabetic Mice. Diabetes. 2016;65(12):3718–29.

27. Becker RE, Berube BJ, Sampedro GR, DeDent AC, and Wardenburg JB. Tissue-specific patterning of host innate immune responses by Staphylococcus aureus α-toxin. Journal of innate immunity. 2014;6(5):619–31.

28. Novelli M, Savoia P, Cambieri I, Ponti R, Comessatti A, Lisa F, et al. Collagenase digestion and mechanical disaggregation as a method to extract and immunophenotype tumour lymphocytes in cutaneous T‐cell lymphomas: Experimental dermatology• Original article. Clinical and Experimental Dermatology: Experimental dermatology. 2000;25(5):423–31.

29. Filgueiras LR, Brandt SL, Wang S, Wang Z, Morris DL, Evans-Molina C, et al. Leukotriene B4-mediated sterile inflammation promotes susceptibility to sepsis in a mouse model of type 1 diabetes. Sci Signal. 2015;8(361):ra10.

30. Bolte S, and Cordelières FP. A guided tour into subcellular colocalization analysis in light microscopy. Journal of microscopy. 2006;224(3):213–32.

31. Abràmoff MD, Magalhães PJ, and Ram SJ. Image processing with ImageJ. Biophotonics international. 2004;11(7):36–42.

32. Huang L, Zhao A, Wong F, Ayala JM, Struthers M, Ujjainwalla F, et al. Leukotriene B4 strongly increases monocyte chemoattractant protein-1 in human monocytes. Arterioscler Thromb Vasc Biol. 2004;24(10):1783–8.

33. Saiwai H, Ohkawa Y, Yamada H, Kumamaru H, Harada A, Okano H, et al. The LTB4-BLT1 axis mediates neutrophil infiltration and secondary injury in experimental spinal cord injury. Am J Pathol. 2010;176(5):2352–66.

34. Netea MG, van de Veerdonk FL, van der Meer JW, Dinarello CA, and Joosten LA. Inflammasome-independent regulation of IL-1-family cytokines. Annu Rev Immunol. 2015;33:49–77.

35. Yang Y, Wang H, Kouadir M, Song H, and Shi F. Recent advances in the mechanisms of NLRP3 inflammasome activation and its inhibitors. Cell death & disease. 2019;10(2):1–11.

36. Brandt SL, and Serezani CH. Too much of a good thing: How modulating LTB4 actions restore host defense in homeostasis or disease. Semin Immunol. 2017;33:37–43.

37. Yokomizo T, Izumi T, Chang K, Takuwa Y, and Shimizu T. A G-protein-coupled receptor for leukotriene B 4 that mediates chemotaxis. Nature. 1997;387(6633):620–4.

38. Afonso PV, Janka-Junttila M, Lee YJ, McCann CP, Oliver CM, Aamer KA, et al. LTB4 is a signal-relay molecule during neutrophil chemotaxis. Developmental cell. 2012;22(5):1079–91.

39. Rogerio AP, and Anibal FF. Role of leukotrienes on protozoan and helminth infections. Mediators of inflammation. 2012;2012.

40. Peters-Golden M, and Henderson WR, Jr. Leukotrienes. N Engl J Med. 2007;357(18):1841–54.

41. Tobin DM, Roca FJ, Oh SF, McFarland R, Vickery TW, Ray JP, et al. Host genotype-specific therapies can optimize the inflammatory response to mycobacterial infections. Cell. 2012;148(3):434–46.

42. Tobin DM, Vary JC, Jr., Ray JP, Walsh GS, Dunstan SJ, Bang ND, et al. The lta4h locus modulates susceptibility to mycobacterial infection in zebrafish and humans. Cell. 2010;140(5):717–30.

43. Brandt SL, Wang S, Dejani NN, Klopfenstein N, Winfree S, Filgueiras L, et al. Excessive localized leukotriene B4 levels dictate poor skin host defense in diabetic mice. JCI Insight. 2018;3(17).

44. Chaves MM, Sinflorio DA, Thorstenberg ML, Martins MDA, Moreira-Souza ACA, Rangel TP, et al. Non-canonical NLRP3 inflammasome activation and IL-1β signaling are necessary to L. amazonensis control mediated by P2X7 receptor and leukotriene B4. PLoS pathogens. 2019;15(6):e1007887.

45. Serezani CH, Aronoff DM, Jancar S, Mancuso P, and Peters-Golden M. Leukotrienes enhance the bactericidal activity of alveolar macrophages against Klebsiella pneumoniae through the activation of NADPH oxidase. Blood. 2005;106(3):1067–75.

46. Gaudreault E, Thompson C, Stankova J, and Rola-Pleszczynski M. Involvement of BLT1 endocytosis and Yes kinase activation in leukotriene B4-induced neutrophil degranulation. The Journal of Immunology. 2005;174(6):3617–25.

47. Campos M, Serezani C, Peters-Golden M, and Jancar S. Differential kinase requirement for enhancement of FcγR-mediated phagocytosis in alveolar macrophages by leukotriene B4 vs. D4. Molecular immunology. 2009;46(6):1204–11.

48. Chung I-C, Yuan S-N, OuYang C-N, Lin H-C, Huang K-Y, Chen Y-J, et al. Src-family kinase-Cbl axis negatively regulates NLRP3 inflammasome activation. Cell death & disease. 2018;9(11):1–14.

49. Sánchez-Galán E, Gómez-Hernández A, Vidal C, Martín-Ventura JL, Blanco-Colio LM, Muñoz-García B, et al. Leukotriene B4 enhances the activity of nuclear factor-κB pathway through BLT1 and BLT2 receptors in atherosclerosis. Cardiovascular research. 2009;81(1):216–25.

50. Ito M, Shichita T, Okada M, Komine R, Noguchi Y, Yoshimura A, et al. Bruton’s tyrosine kinase is essential for NLRP3 inflammasome activation and contributes to ischaemic brain injury. Nature communications. 2015;6:7360.

51. Wang Y, Tao J, and Yao Y. Prostaglandin E2 activates NLRP3 inflammasome in endothelial cells to promote diabetic retinopathy. Hormone and Metabolic Research. 2018;50(09):704–10.

52. Tager AM, and Luster AD. BLT1 and BLT2: the leukotriene B4 receptors. Prostaglandins, leukotrienes and essential fatty acids. 2003;69(2-3):123–34.

53. Serezani CH, Aronoff DM, Sitrin RG, and Peters-Golden M. FcγRI ligation leads to a complex with BLT1 in lipid rafts that enhances rat lung macrophage antimicrobial functions. Blood, The Journal of the American Society of Hematology. 2009;114(15):3316–24.

54. Whang JA, and Chang BY. Bruton’s tyrosine kinase inhibitors for the treatment of rheumatoid arthritis. Drug discovery today. 2014;19(8):1200–4.

55. Lee K-G, Xu S, Kang Z-H, Huo J, Huang M, Liu D, et al. Bruton’s tyrosine kinase phosphorylates Toll-like receptor 3 to initiate antiviral response. Proceedings of the National Academy of Sciences. 2012;109(15):5791–6.

56. Weber AN, Bittner Z, Liu X, Dang T-M, Radsak MP, and Brunner C. Bruton’s tyrosine kinase: an emerging key player in innate immunity. Frontiers in immunology. 2017;8:1454.

57. Park H, Wahl MI, Afar DE, Turck CW, Rawlings DJ, Tam C, et al. Regulation of Btk function by a major autophosphorylation site within the SH3 domain. Immunity. 1996;4(5):515–25.

58. Cho JS, Guo Y, Ramos RI, Hebroni F, Plaisier SB, Xuan C, et al. Neutrophil-derived IL-1β is sufficient for abscess formation in immunity against Staphylococcus aureus in mice. PLoS pathogens. 2012;8(11).

59. Dennis EA, and Norris PC. Eicosanoid storm in infection and inflammation. Nature Reviews Immunology. 2015;15(8):511–23.

