## Supplementary Figures for "Leukotriene B_4_ licenses inflammasome activation to enhance skin host defense"

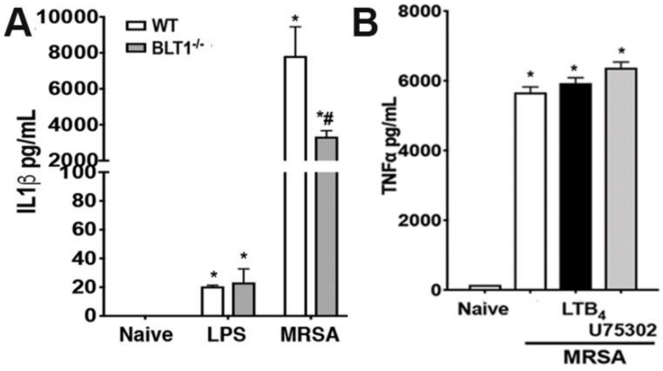


**Supplementary Figure 1. BLT1 receptor enhances IL-1β, but not TNFα production. A)** IL-1β production of macrophages from WT or BLT1-/- mice stimulated with 100 ng/mL LPS or 10:1 MRSA for 24h. **B)** Macrophages were pretreated with 10nM $LTB_4$ for 5 minutes, 10μM U-75302 for 30 minutes, and cultured with or MRSA for 3h followed by TNF-α supernatant quantification. Data are the mean ± SEM from 3 experiments. * $p < 0.05$ *vs.* naive. #$p < 0.05$ *vs.* WT+MRSA.

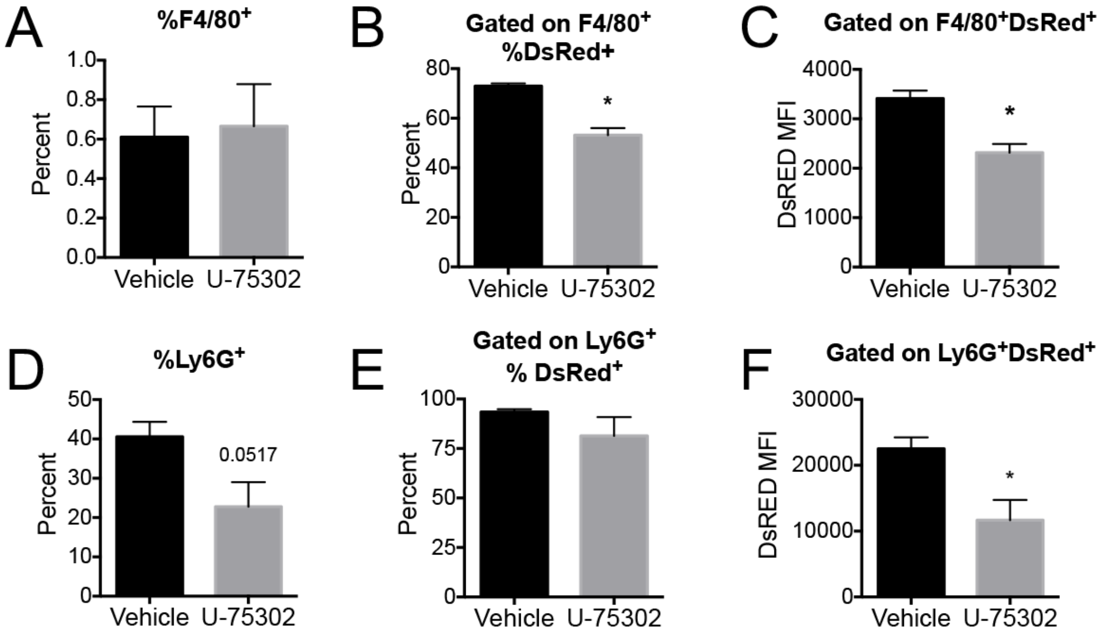


**Supplementary Figure 2. Role of BLT1 in *in vivo* IL-1β expression in macrophages and neutrophils.** pIL1DsRED mice were infected with MRSA and treated topically with vehicle-control or 0.001% BLT1 antagonist U-75302. Biopsy punches were collected at day 1 post-MRSA skin infection and processed for flow cytometry. All cells were gated on, excluding doublets and dead cells for analysis. **A)** Percentage of macrophages (F4/80). **B)** Percentage of F4/80$^+$ macrophages expressing DsRed and **C)** MFI of DsRed. **D)** Percentage of neutrophils (Ly6G). **E)** Percentage of Ly6G$^+$ neutrophils expressing DsRed and **F)** MFI of DsRed. Data are mean ± SEM of 3 mice from 2 experiments. *$p < 0.05$ *vs.* untreated.

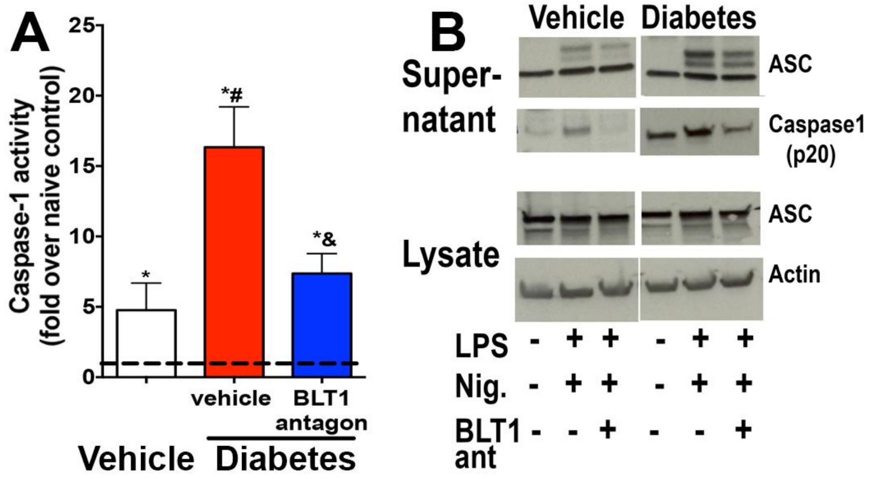


**Supplementary Figure 3. BLT1 receptor drives exaggerated inflammasome activation in macrophages from diabetic mice. A)** Peritoneal macrophages from diabetic and nondiabetic mice were pretreated with 10μM U-75302 (BLT1 antagonist) for 30 min and then challenged with MRSA for 3h. Intracellular active caspase-1 was determined fluorometrically. Data are expressed as the mean ± SEM from 3 individual experiments. *p<0.05 *vs.* vehicle control or WT mice. #p<0.05 *vs.* STZ-treated mice; & p<0.05 *vs.* MRSA. **B)** Macrophages from diabetic and nondiabetic mice were treated with 100ng/mL LPS for 1 h, followed by BLT1 antagonist and 1μM nigericin in the presence of U-75302. Activate caspase-1 and ASC expression were evaluated by immunoblotting. Data are mean of two independent experiments.
